# Flight morphology and visual obstruction predict collision risk in birds

**DOI:** 10.1101/2020.07.20.212985

**Authors:** Erin K. Jackson, Jared A. Elmore, Scott R. Loss, Benjamin M. Winger, Roslyn Dakin

**Affiliations:** Department of Biology, Carleton University, 1125 Colonel By Drive, Ottawa, Ontario, Canada K1S 5B6; Department of Natural Resource Ecology and Management, Oklahoma State University, 008C Ag Hall, Stillwater, OK, USA, 74078; Museum of Zoology, Department of Ecology and Evolutionary Biology, University of Michigan, 1105 North University Avenue, Ann Arbor, MI, USA, 48109

**Keywords:** flight, vision, sensory ecology, mortality, birds, conservation

## Abstract

Collisions with buildings are a major source of mortality for wild birds, but these instantaneous events are difficult to observe. As a result, the mechanistic causes of collision mortality are poorly understood. Here, we evaluate whether sensory and biomechanical traits can explain why some species are more collision-prone than others. We first examined concordance of species vulnerability estimates across previous North American studies to determine whether these estimates are repeatable, and whether vulnerability is more similar among closely-related species. We found moderate concordance and phylogenetic signal, indicating that some bird species are consistently more collision-prone than others. We next tested whether morphological traits related to flight performance and sensory guidance explain these differences among species. Our comparative analysis shows that two traits primarily predict collision vulnerability within passerines: relative beak length and relative wing length. Small passerine species with relatively short wings and those with relatively long beaks are more collision-prone, suggesting that greater maneuverability and obstructed vision contribute to risk. Together, these findings can help inform mitigation strategies and predict which species will be most at risk in other regions.

## 1. INTRODUCTION

Building collisions are the second leading source of direct, human-caused bird mortality in North America, and are estimated to cause between 365–988 million deaths annually in the US [1] and between 16–42 million deaths annually in Canada [2]. Birds are attracted to lighted structures at night [3] or attempt to fly through windows in the daytime [4], only to collide and die upon impact or from injuries later on. Migratory species are especially at risk; the majority of collisions occur during migration, particularly in areas with strong levels of artificial light [3,5]. Additionally, several recent studies have established habitat-related correlates of collision mortality, such as building size and type, glass surface area, light pollution, surrounding land cover, and regional urbanization [6–10]. Behavioural correlates of collisions have also been identified, including nocturnal migration behavior [5,11], insectivory [10,12] and the use of social flight calls [3]. Knowledge of the sensory and biomechanical causes of building collisions is needed to devise effective strategies to prevent these fatalities. However, a major challenge to studying bird collisions with buildings and other anthropogenic structures is that each collision event is instantaneous and difficult to observe and predict.

In the absence of direct observations of collisions, an alternative way to evaluate causes is to consider the physiological traits of species that are most at risk. Not all bird species are equally vulnerable to building collisions; certain species like the white-throated sparrow, black-throated blue warbler, and ruby-throated hummingbird have been found to collide more frequently than expected, given their abundance at the spatial scale over which collisions are counted [1,5,10,11]. Shared traits among these vulnerable species can be a clue to common causes of collisions, and can allow risk to be predicted in other populations that have not been studied [3]. Here, we evaluate whether traits related to sensory guidance and flight performance predict the vulnerability of bird species to collisions. As a first step, we analyzed the concordance among published species vulnerability estimates from four previous studies of North American birds [1,3,10,11]. These four studies are based on >106,000 bird fatalities across 62 geographic locations, and focus primarily on large buildings, with substantial variation among studies in terms of landscape type, building type, survey methods, analytical methods, the spatial scale of the analysis, and site-specific environmental features, such as vegetation and artificial light (see Table S1 for details). This variation is important, because it allows us to identify whether there are repeatable differences in collision risk that are detected across methodologies. If species traits contribute to collision risk, we expect vulnerability estimates to be both repeatable across different studies, and also more similar among closely-related species, as found in a previous study of a single metropolitan region [11].

Next, we evaluated whether species risk is related to traits that influence vision, sensory processing, and flight maneuverability, all of which are expected to influence threat detection and avoidance in flight. Our predictions are explained below and listed in (Table 1). Note that sensory and biomechanical traits may also be correlated with broad classes of foraging and migration behavior, which have been examined in other recent work [7,10]. Our goal is not to address these ecological factors, but rather we aim to identify which (if any) of the sensory and biomechanical traits in Table 1 contribute to collision risk. This is important for a fundamental understanding of bird flight as well as identifying the proximate causes of building collisions.

**Table 1.** Traits that are expected to influence collision vulnerability. In most cases, it is the relative size of the trait (adjusted for body mass) that affects flight. The predicted effect is the expected change in vulnerability when the trait value increases.

| Trait | Relevance for flight | Predicted effect on collision vulnerability |
| --- | --- | --- |
| Absolute eye size | Visual acuity | Decrease |
| Relative eye size | Visual acuity | Decrease |
| Relative beak width | Visual obstruction | Increase |
| Relative beak length | Visual obstruction | Increase |
| Relative skull size | Sensory processing | Decrease |
| Relative wing length | Maneuverability | Increase |
| Relative keel size | Maneuverability | Decrease |

Vision is the primary sense that guides bird flight. Hence, the first set of traits we considered relate to visual guidance. Larger eyes permit greater sensitivity and acuity of vision [13], since they can hold more retinal photoreceptors and receive more light through a larger pupil [14]. We therefore hypothesized that eye size may influence collision risk, and we predicted that birds with larger eyes, and thus greater visual acuity, may have lower vulnerability scores. Another trait that impacts frontal vision is the beak; long beaks relative to body size can produce a longer anterior blind area, and consequently, a narrower binocular field [15]. This blind area may limit the ability for birds to detect oncoming objects or to gauge distance, particularly when objects lie in the frontal plane. Thus, we predicted that species with larger beaks may be at greater risk of colliding. We also predicted that species with enhanced visual processing due to a larger brain [16] may have lower vulnerability scores, since visual processing ability could aid birds in detecting an imminent collision. We expect that these predictions would generally apply to diurnal, nocturnal, and artificial light-induced collisions, because birds with greater visual acuity and less obstructed vision should be better able to correct for visual guidance errors across contexts.

The second set of traits we considered are related to a bird’s maneuverability, defined as the ability to change speed and direction [17]. The ability to generate power to perform rapid evasive maneuvers may allow a bird to avoid collisions more easily. Since enhanced maneuverability is associated with increased flight muscle capacity [18,19], we predicted that birds with greater pectoral muscle size may be less prone to colliding. Differences in maneuverability across bird species are also influenced by wing shape and aspect ratio, such that short broad wings are associated with increased power [20]. Variation in wing shape within a population has also been shown to predict mortality due to vehicle collisions. For example, in cliff swallows, road-killed individuals were found to have significantly longer wings than average in the population, indicating that shorter wings provide an advantage when avoiding vehicle collisions [21]. Thus, we predicted that species with shorter wing spans would have lower collision risk.

Although we generally predicted that species with greater visual and flight capacity would be less collision-prone (Table 1), an alternative scenario is that they may actually be more prone to colliding. For example, highly maneuverable birds with shorter wingspans often fly in more cluttered environments [22], and consequently they may have narrower safety margins that cause them to fly closer to buildings, putting themselves at greater risk. Consistent with this scenario, bird species with greater aerial maneuverability have recently been found to collide more often with aircraft [23]. Similarly, it is possible that species with greater visual capability have narrower safety margins, and that this could lead to elevated risk. Finally, as outlined in Table 1, it is the relative size of most traits (adjusted for body mass) that affects flight, and hence we considered mass-adjusted traits in our analysis. Body mass itself may also contribute to collision risk because it can affect maneuverability, safety margins, and the force of impact during collisions. Therefore, we considered the possibility that differences in body mass may interact with the traits listed in Table 1 to explain why some species collide more than others.

## 2. MATERIAL AND METHODS

### (a) Species collision vulnerability

We compiled estimates of species collision vulnerability from four previous studies that surveyed bird collision fatalities in North America, and that focused primarily on large buildings during spring and fall migration seasons [1,3,10,11] (see Table S1 for details of each study). Two of the studies were conducted at a continent-wide scale, while the other two focused on single metropolitan areas; all four studies differed substantially in monitoring protocol and coverage. Each study reported species vulnerability estimates based on an analysis of collision frequency relative to abundance at the spatial scale over which collisions were counted. Hence, a species reported to have high vulnerability is one that was found to have died in collisions near buildings more often than expected, given its abundance. In total, we gathered 397 vulnerability estimates for 164 species (128 passerines with 2.7 estimates per species on average, and 36 non-passerines with 1.6 estimates per species). We standardized the estimates by scaling each study to have a mean vulnerability of 0 and a standard deviation of 1, to facilitate comparison across studies. A complete list of all species and their standardized vulnerability estimates is provided in the supplement.

We analyzed the concordance of the standardized vulnerability estimates by examining Pearson’s correlations and repeatability using R 3.6.2 [24]. To estimate repeatability, we fit a mixed-effects model of the estimates with species as a random effect and study (four levels) as the fixed effect in the lme4 1.1-21 package [25], and we calculated repeatability as the proportion of variance attributed to species ID [26]. We used parametric bootstrapping to obtain the 95% CIs for repeatability. To determine whether vulnerability is more similar among closely related species, we first calculated species-average vulnerability by taking the mean of the available standardized estimates (note that if a species was not common at a particular study’s geographic location, it would not have an estimate). We used the phylogeny from http://BirdTree.org [27,28] to calculate phylogenetic lambda of species-average vulnerability scores using the phytools 0.6-99 package [29]. Lambda is a measure of phylogenetic signal that ranges from zero to one. A lambda value near one indicates that vulnerability is highly similar for closely related species, whereas a value near zero indicates that vulnerability is independent of phylogenetic relatedness.

### (b) Morphological measurements

We collected data on morphological traits representative of each species from skeletal specimens available at the Canadian Museum of Nature. We aimed to measure four to five adult-sized individuals per species, including both sexes, where available. Our measurements included 393 specimens from 89 species (4.4 ± SD 1.2 individuals per species), with variation due to the number of intact specimens in the collection. Most of the studied species breed across the North American continent (see supplement for details). For each individual specimen, we recorded body mass from the specimen metadata where available, and photographed the scleral rings, the skull from the dorsal view, and the humerus and ulna using a digital SLR camera (Canon EOS M50 with EF-M 28 mm lens) mounted on a copy stand. As an indication of flight muscle capacity, we measured the keel along the diagonal axis from the anterior ventral point to the posterior ventral point using digital calipers (± 0.01 mm). The length of the keel is highly correlated with the mass of the primary flight muscles [30,31], which provide the main source of power for flight maneuvers [19].

We measured eye size, beak length and width, skull size, and wing length from digital images using ImageJ 1.52a [32]. All pixel measurements in ImageJ were converted to cm using a 5 cm scale that was included in every photograph. As a measure of eye size, we measured the inner diameter of each scleral ring – a ring of overlapping bony plates inside the sclera of the eye [33]. Scleral ring diameter is highly correlated with eye size and provides a proxy of spatial visual acuity [34]. For any specimen with both scleral rings intact, we averaged the left and right diameters. As a measure of visual obstruction, we measured the length and width of the beak from the dorsal view of the skull. Beak width was measured at the base of the beak where it meets the skull, and beak length was measured as the length of the skull plus the beak. As a proxy for brain size variation across species, we measured the width of the skull at its widest point. External head size and skull volume have been shown to reliably predict inter-individual variation in brain mass [35,36]. Hence, the width of the skull is expected to be a reliable measure of species differences in our sample. As our measure of wing length, we took the length of each humerus and ulna as the longest straight-line distance (chord). For the humerus, this measurement extended from the tip of the shoulder joint to the inside of the elbow notch. If an individual had two matching bones intact (e.g., left and right humerus), we averaged the two measures. Some specimens (and species) were missing traits (e.g., due to broken bones); details of the sample sizes per trait are provided in Fig. S1.

For each of the traits described above, we computed the species-average trait value relative to body mass, according to our hypotheses in Table 1. This is important because the species studied vary widely in body mass (Fig. S1), and the capacity of a given anatomical trait to influence flight also depends on body mass. To compute each relative trait value, we first fit a linear model of the log-transformed trait values relative to log-transformed body mass, across all individual specimens (Fig. S1), and extracted the model residuals. For the specimens that did not have body mass available in the metadata (26% of 393 individuals), we used the mean species mass for adults from the Birds of the World online database [37]. Finally, we took the mean value of the residuals for each species as the relative species trait value (see the supplement for a detailed explanation). Hence, a species with a large relative keel size has a larger keel than expected for its body mass. The only trait where we also considered absolute magnitude was eye size (Table 1).

Because relative humerus length was closely correlated with relative ulna length (r = 0.81, p < 0.0001, n = 88 species), we combined these two traits into one measure of species relative wing length by taking the average. We also verified that both humerus and ulna length are highly correlated with species wing chord, a common measure of the distal part of the wing (r = 0.90 and 0.92 respectively, all p < 0.0001, see supplement for details). Prior to further analysis, we checked variance inflation factors for all of the relative trait values and body mass, and confirmed that the variance inflation factors were relatively low (all < 2.3).

### (c) Collision vulnerability in relation to morphological traits

To evaluate whether visual and flight-related traits predict collision vulnerability, we modelled the standardized vulnerability estimates using Bayesian phylogenetic mixed-effects regression in the package MCMCglmm 2.29 [38]. The models included a random effect of species to account for repeated measures, as well as the phylogenetic structure to account for non-independence among related species [19,39]. The use of a repeated measures analysis ensured that species with more estimates were accorded greater weight in the analysis. We fit a series of candidate models to consider hypothesized effects of sensory and biomechanical traits; these candidate models are listed in Table S2. Each candidate model included fixed effects of either one, two, or none of the skeletal traits listed in Table 1. Additional fixed effects accounted for the source study (categorical variable with four levels), a species’ log-transformed body mass (continuous variable), and a binary factor describing whether a species was a nocturnal migrant that used a social flight call. Previous evidence has shown that nocturnal migration with social flight call use is an important predictor of urban collision risk [3].

We built two model sets. The “All birds” model set analyzed passerines and non-passerines together (n = 205 vulnerability estimates for 85 species), whereas the “Passerine only” analysis was limited to the order Passeriformes (n = 164 vulnerability estimates for 59 species). There were no non-passerines < 40 g in our morphometric analysis, and only two passerines > 100 g. This makes it difficult to distinguish effects that are due to small size vs. membership in the passerine clade. Hence, running both model sets provided a check on whether our conclusions were robust to taxonomic and size-based sampling. The “All birds” models included additional fixed effects for membership in the passerine order and the potential interaction between each trait and body mass (i.e., size-dependence of trait effects). Because we used model comparison, the analyses only included species for which complete trait data were available. We used DIC (deviance information criterion) to compare Bayesian models fit with MCMC.

Each model was run for 500,000 iterations after a burn-in period of 3,000, and posterior results were thinned to include every 500^th^ sample. We verified convergence using the Gelman-Rubin statistic and confirmed that posterior autocorrelation values were < 0.1. As a means of visualizing the main results, we refit the top models from the MCMCglmm analysis with lme4 1.1-21 [25] using taxonomic family and species as nested random effects. After confirming that the coefficients from this lme4 analysis were consistent with the posterior MCMCglmm model, we plotted the results using partial residual plots from the visreg 2.5-0 package [40].

## 3. RESULTS

### (a) Consistency and phylogenetic distribution of collision risk

Published estimates of bird species collision vulnerability were moderately repeatable (R = 0.32 [95% CI 0.20 to 0.43]) and positively correlated (average r of 0.33; Fig. 1a). Examining the composite vulnerability score (all studies combined), we find that closely related species tend to have similar collision risk (Fig. 1b), with a moderate phylogenetic signal (lambda = 0.54, p < 0.001).

**Figure 1.**
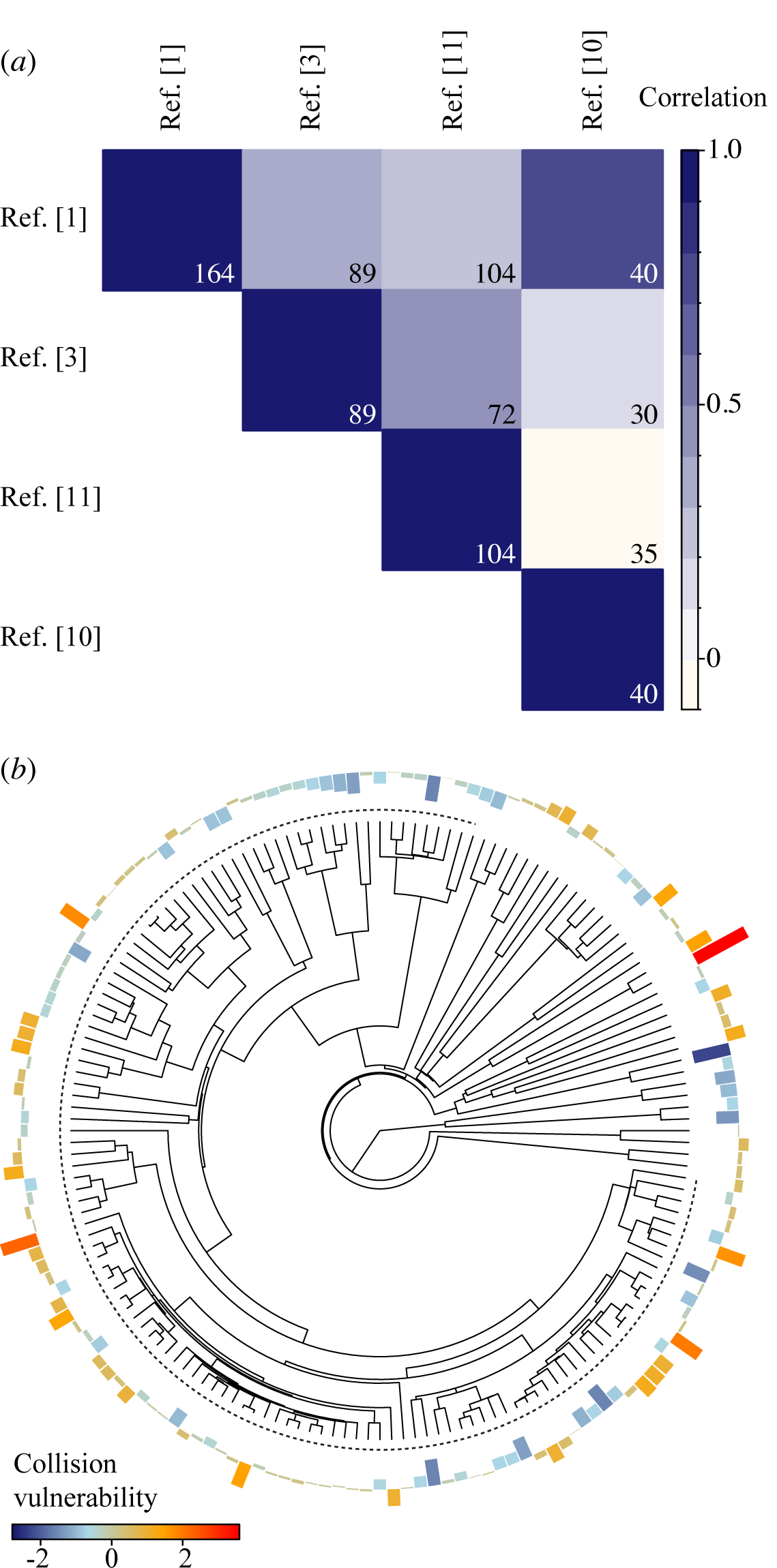
Estimates of bird species vulnerability to collisions are repeatable. (a) The heatmap shows Pearson’s correlation coefficients comparing estimates from four previous studies. Most correlations were positive and moderately strong. Sample sizes (i.e., numbers of species) are shown within each cell; see the list of References [#] and Table S1 for details. (b) Distribution of species-average standardized collision vulnerability across the avian phylogeny (n = 164 species). Dark blue indicates species with low composite vulnerability (colliding less often than expected, based on abundance), whereas red indicates high vulnerability. The dotted line denotes the passerines. See Fig. S2 for additional taxonomic information.

### (b) Collision vulnerability and morphology

The best-fit models of species collision vulnerability indicated that two skeletal traits were associated with increased risk: relatively long beaks and relatively short wings (Fig. 2; Tables S3-S4). In the “All birds” analysis, the effect of beak and wing length was limited to small and medium-sized species (Table S5), which are almost exclusively the passerines in our dataset. The best-fit “Passerine only” model led to the same conclusion. Both model sets also identified body mass as an important predictor, with smaller species generally being more collision prone (Fig. S3 and Tables S4-S5). Consistent with a previous study [3], collision risk was also found to be higher for nocturnal migrants that use social flight calls (Fig. S3).

**Figure 2.**
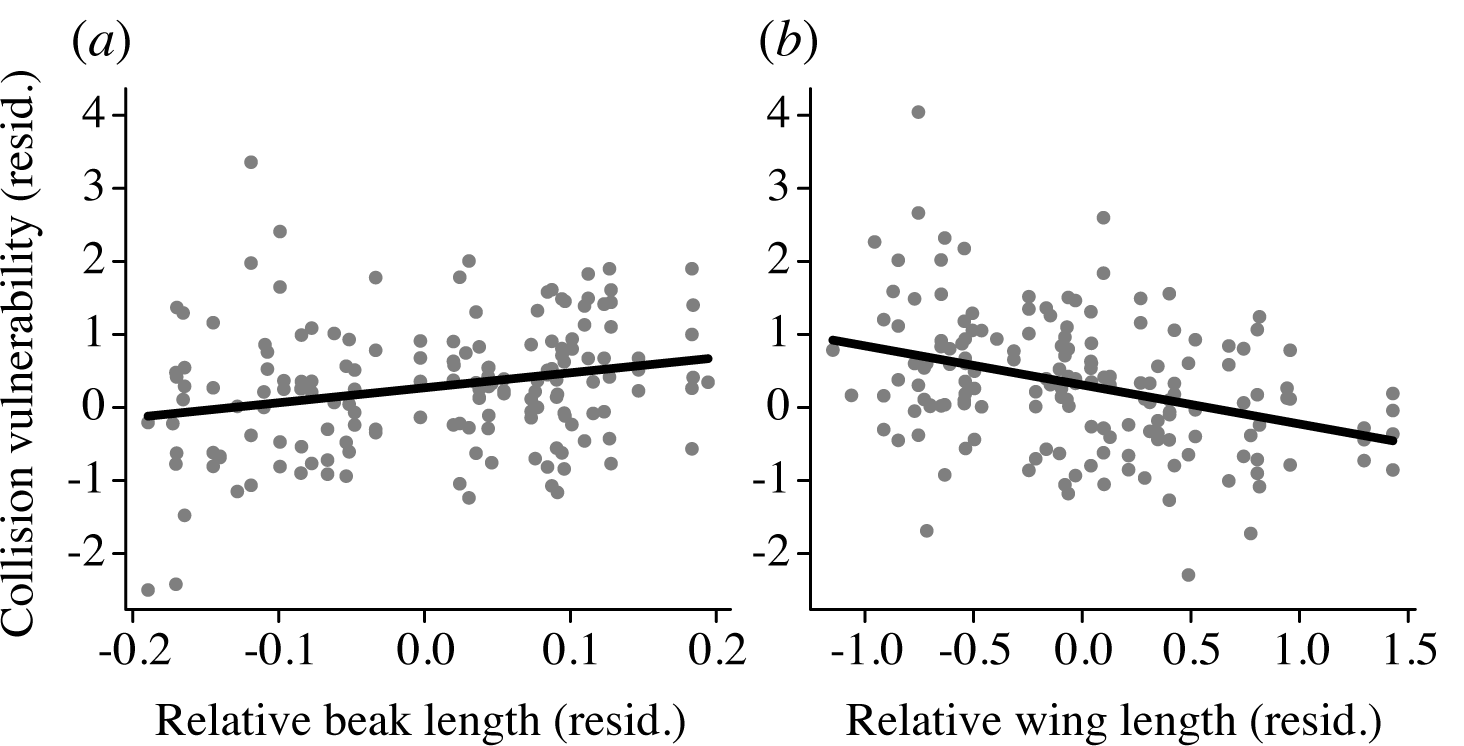
Collision risk for small passerines is associated with maneuverability and visual obstruction. Partial residual plots show the relationship between collision vulnerability and morphological traits that were retained in the best-fit model (“Passerine only”). Each data point represents a residual vulnerability estimate after accounting for the effects of nocturnal migration calling behaviour and body mass (n = 59 species; see Fig. S3 for additional data). Within passerines, relatively elongated beaks (a) and short wings (b) are associated with elevated rates of collisions. The full results of this analysis are provided in the supplement. Beak and wing length are also found to predict risk for small species in the “All birds” model (Table S5).

## 4. DISCUSSION

Several recent studies have established that bird species differ in vulnerability to building collisions [1,5,10,11]. We show here that despite pronounced differences across studies in terms of geographic scale and monitoring protocols, estimates of species vulnerability are moderately repeatable and associated with phylogeny (Fig. 1).

In addition to factors that have previously been studied [3,10,11], our findings indicate that visual obstruction may be one of the contributing factors to collision risk. Vulnerability was positively associated with beak length for small passerine birds (Fig. 2a). Examples of species with relatively long beaks that are consistently identified as collision-prone include the white-breasted nuthatch, brown creeper, and gray catbird (Table S7). This general result suggests that long beaks that obstruct the frontal visual field [15] may also impair the ability of small birds to avoid collisions. With impaired vision, a bird might be more apt to misinterpret window reflections as vegetation or open sky, or be more prone to guidance errors caused by artificial light. Smaller birds also have visual systems with lower spatial resolution [14,41,42], and a long beak obstructing their frontal visual field might exacerbate this issue.

Our results point to maneuverability as another contributing factor to collision risk, because small passerine species with relatively short wings were found to be more collision-prone (Fig. 2b). Examples of short-winged species include the white-throated sparrow, fox sparrow, and rose-breasted grosbeak, all of which are estimated to have high collision risk (Table S7). Given that short, broad wings allow for increased power and maneuverability [20], this finding suggests that more maneuverable species are more collision-prone. This result also contradicts our initial prediction in Table 1, but is consistent with another recent study finding that more maneuverable bird species are more likely to strike aircraft [23]. Highly maneuverable species often fly in cluttered environments such as forest and dense vegetation [22]. They may thus have narrower safety margins that cause them to venture closer to buildings in urban and suburban environments, both during the day and at night when errors may be caused by artificial light.

Our results indicate that small body size is generally associated with increased risk, even within passerines (Fig. S3). We propose that this is because small body size is also associated with narrow safety margins during flight. Small birds are often observed to enter denser vegetation and other cluttered airspaces as an escape strategy (e.g., when chased by predators and/or competitors) [43]. They may also depend on high-risk airspaces for foraging opportunities. Hence, buildings and window barriers may create ecological traps for the smallest and most maneuverable bird species. In line with our general results, hummingbirds are estimated to be the most collision-prone group of bird species [1], and they also have extremely small body sizes, high maneuverability, and long beaks. Although we were not able to collect morphometric data from hummingbirds, our results point to their common traits as generally collision prone.

There are many other traits relevant to flight control that should be investigated in future collision studies. For example, not all birds look forward when they fly [44]. Collision avoidance may thus depend on the orientation of the eyes as well as the position of the eyes relative to the beak [45]. Similarly, the temporal resolution of vision (number of changes perceived per second) is another important trait that may influence the ability of a bird to rapidly detect and avoid obstacles [46]. It is important to note that a vast number of collisions – particularly among passerines – occur during nocturnal and crepuscular times when birds can be disoriented or attracted by artificial light sources [3]. Monitoring studies that examine collisions at those times can help determine whether certain visual systems are more sensitive to guidance errors caused by artificial light, and how we might prevent bird strikes under these conditions.

Previous studies have provided great insight into the features of urban and suburban environments that pose risk, such as artificial light at night, glass surface area, and proximity to vegetation [3,6–8,11,12]. Our results show how morphology can also explain why some species are more collision-prone than others, owing to their small size, visual obstruction, and maneuverability. Our findings also have implications for predicting the impacts of building development on bird populations outside of North America, where monitoring data may be deficient. Mitigation strategies are often costly, and understanding which morphological and flight-related traits lead to collisions will help target these strategies towards a particular trait or species group. For example, it will be important to ensure that the placement, spacing, colouration, and size of features used in products designed to make glass more bird-friendly are large and frequent enough that small birds with obstructed vision can detect them [47].

An important caveat is that our results are correlative, and a complete understanding of the proximate causes of collisions still requires directly observing these events. Additionally, nearly all of the fatalities used in our analysis occurred in urban settings during migration. Although urban collisions are better sampled, residential buildings in suburban habitats are estimated to cause a substantial proportion of total bird collision mortality across the continent [1,2,48]. Residential buildings may elicit different visual guidance errors and affect different species than urban and campus buildings [1]. There is an urgent need for studies of the proximate mechanisms of non-fatal collisions, and collisions in suburban environments. In the future, our results suggest that experimental tests of mitigation strategies should be targeted towards small, maneuverable species with obstructed vision in order to have the greatest benefit.

## DATA ACCESSIBILITY

All data and R scripts are available at: https://figshare.com/s/0de0dc5ff609a2d19c15 The repository will be made public when the final version of the study is published.

## AUTHOR CONTRIBUTIONS

RD and EJ conceived the study, collected the data, analyzed the data, and wrote the manuscript. JAE, SRL, and BMW contributed vulnerability estimates and edited the manuscript.

## ACKNOWLEDGEMENTS

This work was possible thanks to many researchers who led collision surveys, as well as hundreds of citizen-scientists and students who contributed to those efforts. We thank the Canadian Museum of Nature, Kamal Khidas, Greg Rand, Kyna Crumley, Kara Scott, Mahmoud El-Saadi, Courtney Donkersteeg, Paisley Clunis, Santiago Claramunt, and Natalie Wright.

