## Supplementary Information for "Flight morphology and visual obstruction predict collision risk in birds"

**Table S1. Sources of collision vulnerability estimates.** The four studies below provided estimates of species collision vulnerability based on the frequency of collision records relative to abundance. The numbers in square brackets [#] refer to the Reference list provided in the main text. Note that the broad analysis by Loss et al. 2014 [1] included a portion of data that overlapped with Nichols et al. 2018 [11] and another portion of data that overlapped with Winger et al. 2019 [3]. However, given substantial differences in methodology, and numerous additional data sources in [1] comprising different locations, building types, and species, we consider these three studies to be distinct estimates. We did not analyze estimates from an earlier study by Arnold and Zink 2011 [5], because it used the same analytical approach as Loss et al., but only a subset of the data in the Loss et al. study.

| | [1] Loss et al. 2014 | [11] Nichols et al. 2018 | [3] Winger et al. 2019 | [10] Elmore et al. 2020 |
|---|---|---|---|---|
| Type of study | Meta-analytic compilation of 23 studies | Systematic monitoring by citizen scientists | Combination of systematic monitoring at a convention centre and monitoring by citizen scientists | Systematic and coordinated monitoring at 40 campuses using a standardized protocol |
| Location(s) used to estimate vulnerability | Multiple sites in USA and Canada | Minneapolis and St. Paul, USA | Chicago, USA; note that the paper also reports additional analyses from other locations, but we used the estimates for Chicago | 40 sites in USA, Canada, and Mexico |
| Landscape type(s) | Urban, rural, residences, academic campuses | Urban | Urban | Academic campuses |
| Description of building type(s) | Low-rise buildings (4 – 11 storey), high-rise buildings (> 11 storey), and residences (1 – 3 storeys); we used the estimates provided for “All buildings” | 2 – 57 storey buildings | Mix of low-rise and high-rise buildings throughout Chicago | 1–14 storey buildings |
| Data collection years | 1969–2012 | 2007–2010 | 1978–2016 | 2014 |
| Times of day | Combination of surveys and opportunistic collection (various times of day) | Morning (0600 – 1000 h) | Combination of predawn surveys and opportunistic collection (various times of day) | Afternoon (1400 – 1600 h) |

| …cont. | [1] Loss et al. 2014 | [11] Nichols et al. 2018 | [3] Winger et al. 2019 | [10] Elmore et al. 2020 |
|---|---|---|---|---|
| Season(s) | Mainly spring and fall migration | Spring and fall migration | Spring and fall migration | Fall migration |
| Method of estimating species vulnerability | Residuals based on frequency of collisions within a given species' range, accounting for abundance | Random effect estimates (intercepts) from a mixed-effects model of weekly collision frequency, accounting for abundance as a fixed effect | Chi-square residual for observed collision frequency vs. expected frequency, based on abundance | Residuals based on frequency of collisions per unit of survey effort within a given species' range, accounting for abundance |
| Source of abundance data | Partners in Flight, continent-wide | Systematic point counts, local abundance | eBird, local abundance | Partners in Flight, continent-wide |
| Includes estimates for resident/migratory species? | Both | Migratory only | Migratory only | Both |
| Includes estimates for non-passerines? | 128 passerines<br>36 non-passerines | 87 passerines<br>17 non-passerines | 89 passerines<br>0 non-passerines | 37 passerines<br>3 non-passerines |

**Adjusting traits for body mass**

It was important to adjust traits for body mass for two reasons. First, the traits listed in Table 1 are expected to influence flight relative to body mass and size. An example is the keel; on its own, a bird species' keel size (and pectoral muscle size) is not informative of its flight performance. Instead, the power available for flight is determined by a bird's keel size relative to its body mass. This is illustrated by hummingbirds, a group of birds noted for their extreme flight agility. Hummingbirds have very small keels in absolute terms, but they are able to generate extreme power during their flight maneuvers because their pectoral muscles (and keel) are greatly hypertrophied. Hence, the relative keel size of hummingbirds is among the largest found in any group of birds.

A second important consideration is that all of the measured traits in our study are highly correlated with inter-species variation in body mass. A regression model with absolute trait values as predictors would have Variance Inflation Factors (VIFs) > 11. This was important because we further wished to account for the potential statistical interaction between body size and mass-adjusted trait values. To resolve this issue, we used the approach described in the main text of first determining mass-adjusted relative trait values at the individual level (per studied museum specimen; using residuals as shown in Fig. S1). Then, we calculated species-averages of these residuals as our "relative" (mass-adjusted) species trait values for further analysis. The advantage of this approach is that it allowed us to model several mass-adjusted traits and body mass as predictors within the same model, while maintaining VIFs < 2.3. Note that when calculating the relative trait values, we did not account for phylogeny, because the trait value is purely a morphological parameter. Phylogeny was accounted for during subsequent statistical inference.

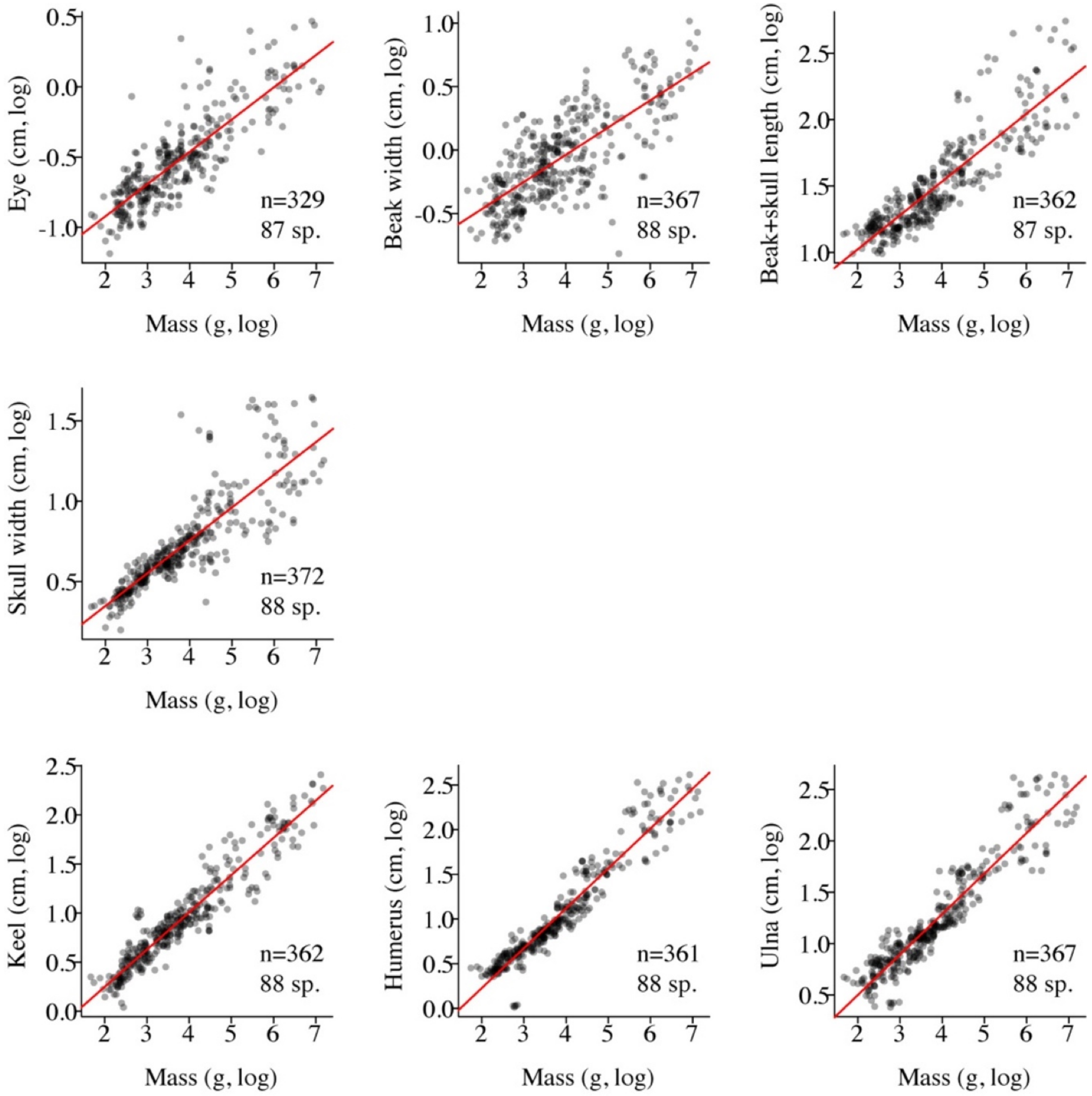

**Figure S1. Individual skeletal traits in relation to body mass.** Data were collected from a total of 393 individual specimens representing 89 North American species. Sample sizes within each panel give the number of individuals (n) and species (sp.). Note that these sample sizes vary because some individuals and species were missing some traits (e.g., due to broken or missing parts). The red lines show the linear regression for the log-transformed trait against log-transformed individual body mass.

**Wing skeletal traits and wing chord**

To check the correspondence between measures of proximal limb bones (humerus and ulna) and typical measures of distal feathered wing length, we downloaded North American avian specimen data from the University of Michigan Museum of Zoology (UMMZ). The UMMZ dataset is available via VertNet http://www.vertnet.org/index.html and includes many entries on wing chord, which is a standard measure of a bird's feathered wing length. We identified all adult wing chord entries that could be matched to one of the 88 North American species in our humerus/ulna dataset (n = 3,853 specimens from the UMMZ collection, representing 85 of the species in our dataset, with n = 45 individuals per species on average). For each species, we computed the average wing chord in the UMMZ dataset. Finally, we quantified the Pearson's correlations between species-average wing chord and species-average humerus and ulna lengths, respectively. The correlation coefficients were consistent within the 95% CI whether we used raw or log-transformed dimensions.

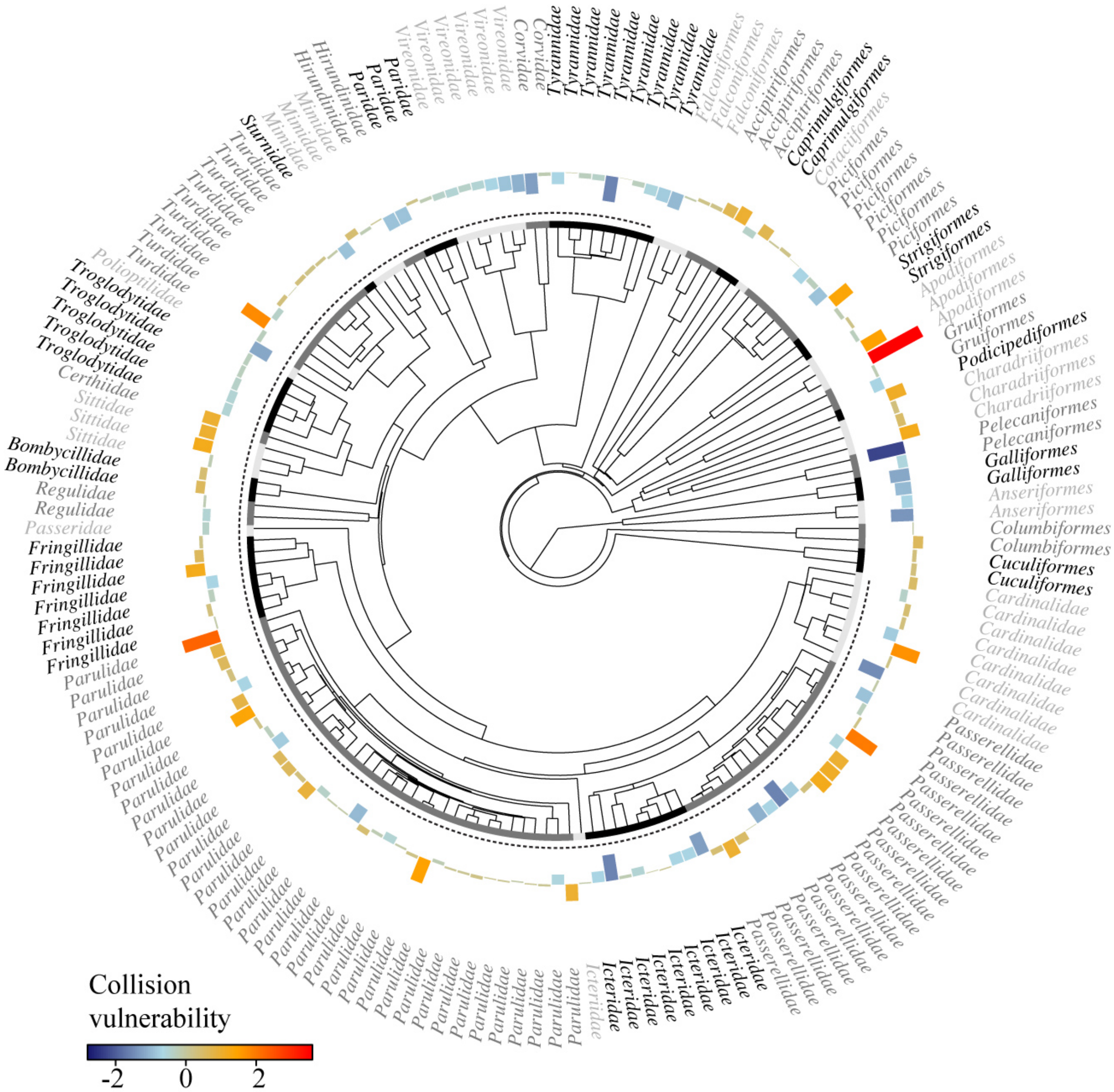

**Figure S2. Phylogenetic distribution of species-average collision vulnerability.** This figure shows the same data illustrated in Fig. 1b, but with additional labels to indicate either taxonomic order (for non-passerines) or family (for the passerine order Passeriformes, which make up the majority of species studied). The shaded bar at the tips of the phylogeny indicates order/family transitions and is shaded to match the labels. The dotted line indicates the passerines.

**Table S2. Candidate models to predict species vulnerability to collisions.** In the notation below, .W and .L refer to width and length, respectively, and .r refers to the relative trait value (i.e., relative to body mass). All the candidate models accounted for study, log-transformed body mass, nocturnal migration flight calls, repeated measures of each species, and the phylogenetic relationships among species. The "All birds" models additionally included a fixed effect for membership in the passerine order and the potential interaction between each trait and body mass. Note that we did not consider any models with both eye.r and eye.cm, because we considered these two measures of eye size to be redundant.

| Model | Skeletal traits |
|---|---|
| 1 | *includes no traits from Table 1* |
| 2 | ~ eye.r |
| 3 | ~ eye.abs |
| 4 | ~ beak.W.r |
| 5 | ~ beak.L.r |
| 6 | ~ skull.W.r |
| 7 | ~ wing.L.r |
| 8 | ~ keel.r |
| 9 | ~ eye.r + beak.W.r |
| 10 | ~ eye.r + beak.L.r |
| 11 | ~ eye.r + skull.W.r |
| 12 | ~ eye.r + wing.L.r |
| 13 | ~ eye.r + keel.r |
| 14 | ~ eye.abs + beak.W.r |
| 15 | ~ eye.abs + beak.L.r |
| 16 | ~ eye.abs + skull.W.r |
| 17 | ~ eye.abs + wing.L.r |
| 18 | ~ eye.abs + keel.r |
| 19 | ~ beak.W.r + beak.L.r |
| 20 | ~ beak.W.r + skull.W.r |
| 21 | ~ beak.W.r + wing.L.r |
| 22 | ~ beak.W.r + keel.r |
| 23 | ~ beak.L.r + skull.W.r |
| 24 | ~ beak.L.r + wing.L.r |
| 25 | ~ beak.L.r + keel.r |
| 26 | ~ skull.W.r + wing.L.r |
| 27 | ~ skull.W.r + keel.r |
| 28 | ~ wing.L.r + keel.r |

**Table S3. The top five best-fit "All birds" models of species collision vulnerability (n = 205 estimates for 85 passerine and non-passerine species).** Models were ranked according to DIC. All models also accounted for study, log-transformed body mass, nocturnal migration flight calls, membership in the passerine order, repeated measures of each species, and the phylogenetic relationships among species.

| Model rank | Skeletal traits | DIC |
|---|---|---|
| 1 | ~ beak.L.r * mass + wing.L.r * mass | 584.3 |
| 2 | ~ wing.L.r * mass | 587.2 |
| 3 | ~ skull.W.r * mass + wing.L.r * mass | 587.6 |
| 4 | *includes no traits from Table 1* | 587.8 |
| 5 | ~ eye.r * mass + wing.L.r * mass | 587.8 |

**Table S4. The top five best-fit "Passerine only" models of species collision vulnerability (n = 164 estimates for 59 passerine species).** Models were ranked according to DIC. All models also accounted for study, log-transformed body mass, nocturnal migration flight calls, repeated measures of each species, and the phylogenetic relationships among species.

| Model rank | Skeletal traits | DIC |
|---|---|---|
| 1 | ~ beak.L.r + wing.L.r | 475.8 |
| 2 | ~ eye.abs + wing.L.r | 476.9 |
| 3 | ~ eye.r + wing.L.r | 477.1 |
| 4 | ~ wing.L.r | 478.1 |
| 5 | ~ eye.r | 478.7 |

**Table S5. Posterior estimates from the best-fit "All birds" model.**

| Fixed effect | Estimate (posterior mean) | Credible interval (95%) | Effective sample size | pMCMC |
|---|---|---|---|---|
| Passerine (y vs. n) | –0.76 | –1.34, –0.21 | 902 | 0.01 |
| Nocturnal mig. w. call (y vs. n) | 0.54 | 0.26, 0.87 | 1,000 | 0.004 |
| Mass (log) | –0.24 | –0.47, –0.01 | 1,000 | 0.04 |
| Beak.L.r | 4.98 | 1.22, 8.48 | 862 | 0.006 |
| Wing.L.r | –0.99 | –1.64, –0.32 | 1,000 | 0.004 |
| Beak.L.r * mass | –1.03 | –1.80, –0.24 | 901 | 0.006 |
| Wing.L.r * mass | 0.18 | 0.06, 0.33 | 1,000 | 0.006 |

**Table S6. Posterior estimates from the best-fit "Passerine only" model.**

| Fixed effect | Estimate (posterior mean) | Credible interval (95%) | Effective sample size | pMCMC |
|---|---|---|---|---|
| Nocturnal mig. w. call (y vs. n) | 0.58 | 0.23, 0.94 | 911 | 0.002 |
| Mass (log) | –0.36 | –0.69, –0.06 | 1,188 | 0.03 |
| Beak.L.r | 2.08 | 0.30, 3.71 | 1,121 | 0.02 |
| Wing.L.r | –0.55 | –0.96, –0.20 | 1,000 | 0.008 |

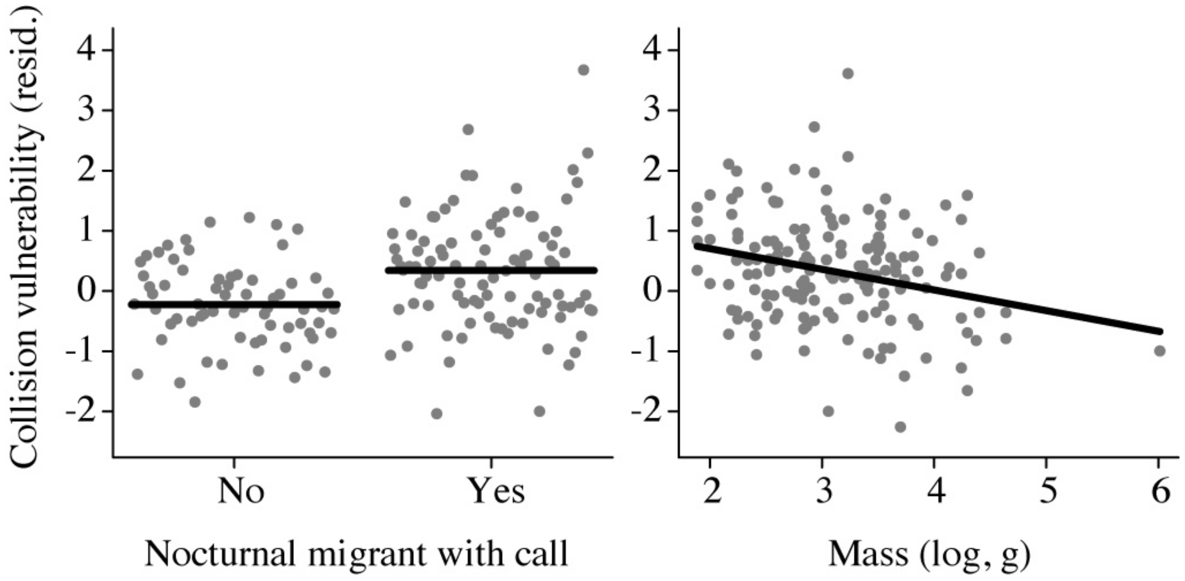

**Figure S3. Collision risk is associated with nocturnal migration call use and small body size within passerines.** See Fig. 2 of the main text for the other predictors from the best-fit model of passerine collision risk (Table S6). Each data point represents a residual estimate after accounting for other factors in the best-fit model. Note that the conclusions of this analysis are unchanged when excluding the largest passerine species, the American crow (*Corvus brachyrhynchos*).

**Table S7. List of species and standardized vulnerability estimates.** For each species, the table lists taxonomic information, the standardized vulnerability estimates in each study, and the percentile vulnerability rank within each study (in bold). Larger positive values indicate that a species is more collision prone. Dashes (--) indicate that a species was not estimated in that study. The table is sorted by taxonomic Order (alphabetically), followed by Family (alphabetically), and then by species-average vulnerability within families (descending). The Morph column indicates whether a species was studied in the statistical analysis of sensory and biomechanical traits. The Breeding column indicates whether a species primarily breeds east of the North American Cordillera ("E and/or C"), west of the North American Cordillera ("Western"), or on both sides of the North American Cordillera ("Cross-cont"). Most of the species in the statistical analysis of sensory/biomechanical traits (65/85) breed across the continent, and most (59/85) are passerines. Shading is used to visually delineate orders among non-passerines, and families within the passerine order. The complete dataset is available in the figshare repository for this study.

| Species | Order | Family | [1] Loss et al. 2014 | | [11] Nichols et al. 2018 | | [3] Winger et al. 2019 | | [10] Elmore et al. 2020 | | Morph | Breeding |
|---|---|---|---|---|---|---|---|---|---|---|---|---|
| *Accipiter striatus* | Accipitriformes | Accipitridae | 0.97 | **84** | -- | **--** | -- | **--** | -- | **--** | Yes | Cross-cont |
| *Accipiter gentilis* | Accipitriformes | Accipitridae | 0.72 | **80** | -- | **--** | -- | **--** | -- | **--** | Yes | Cross-cont |
| *Accipiter cooperii* | Accipitriformes | Accipitridae | 0.78 | **83** | –0.3 | **46** | -- | **--** | -- | **--** | Yes | Cross-cont |
| *Anas platyrhynchos* | Anseriformes | Anatidae | –0.93 | **17** | –0.43 | **35** | -- | **--** | -- | **--** | Yes | Cross-cont |
| *Aix sponsa* | Anseriformes | Anatidae | –2.48 | **2** | –0.31 | **43** | -- | **--** | -- | **--** | Yes | Cross-cont |
| *Chaetura pelagica* | Apodiformes | Apodidae | –0.2 | **39** | –0.31 | **43** | -- | **--** | -- | **--** | -- | E and/or C |
| *Calypte anna* | Apodiformes | Trochilidae | 3.59 | **100** | -- | **--** | -- | **--** | -- | **--** | -- | Western |
| *Archilochus colubris* | Apodiformes | Trochilidae | 1.78 | **99** | 1.19 | **88** | -- | **--** | 1.49 | **95** | -- | E and/or C |
| *Caprimulgus vociferus* | Caprimulgiformes | Caprimulgidae | 0.74 | **82** | -- | **--** | -- | **--** | -- | **--** | -- | E and/or C |
| *Chordeiles minor* | Caprimulgiformes | Caprimulgidae | –0.68 | **23** | –0.2 | **55** | -- | **--** | -- | **--** | Yes | Cross-cont |
| *Charadrius vociferus* | Charadriiformes | Charadriidae | 0.19 | **56** | 0.48 | **77** | -- | **--** | -- | **--** | Yes | Cross-cont |
| *Larus delawarensis* | Charadriiformes | Laridae | 0.48 | **68** | -- | **--** | -- | **--** | -- | **--** | Yes | Cross-cont |
| *Scolopax minor* | Charadriiformes | Scolopacidae | 1.16 | **91** | -- | **--** | -- | **--** | -- | **--** | Yes | E and/or C |
| *Zenaida macroura* | Columbiformes | Columbidae | –0.05 | **46** | 2.79 | **98** | -- | **--** | –0.9 | **20** | Yes | E and/or C |
| *Columba livia* | Columbiformes | Columbidae | –0.06 | **45** | -- | **--** | -- | **--** | -- | **--** | Yes | Cross-cont |

| | | | | | | | | | | | | |
|---|---|---|---|---|---|---|---|---|---|---|---|---|
| *Megaceryle alcyon* | Coraciiformes | Alcedinidae | 0.53 | **71** | –0.34 | **39** | -- | **--** | -- | **--** | -- | Cross-cont |
| *Coccyzus erythropthalmus* | Cuculiformes | Cuculidae | 0.39 | **64** | 0.26 | **71** | -- | **--** | -- | **--** | Yes | E and/or C |
| *Coccyzus americanus* | Cuculiformes | Cuculidae | 0.44 | **66** | 0.07 | **65** | -- | **--** | -- | **--** | Yes | E and/or C |
| *Falco columbarius* | Falconiformes | Falconidae | 0.56 | **73** | –0.16 | **60** | -- | **--** | -- | **--** | Yes | Cross-cont |
| *Falco sparverius* | Falconiformes | Falconidae | –0.01 | **49** | –0.2 | **55** | -- | **--** | -- | **--** | Yes | Cross-cont |
| *Falco peregrinus* | Falconiformes | Falconidae | –1 | **16** | -- | **--** | -- | **--** | -- | **--** | Yes | Cross-cont |
| *Bonasa umbellus* | Galliformes | Phasianidae | –1 | **16** | -- | **--** | -- | **--** | -- | **--** | Yes | Cross-cont |
| *Phasianus colchicus* | Galliformes | Phasianidae | –1.23 | **13** | -- | **--** | -- | **--** | -- | **--** | -- | Cross-cont |
| *Coturnicops noveboracensis* | Gruiformes | Rallidae | –0.18 | **40** | -- | **--** | -- | **--** | -- | **--** | -- | E and/or C |
| *Fulica americana* | Gruiformes | Rallidae | –0.71 | **21** | -- | **--** | -- | **--** | -- | **--** | Yes | Cross-cont |
| *Bombycilla garrulus* | Passeriformes | Bombycillidae | 0.56 | **73** | -- | **--** | -- | **--** | -- | **--** | Yes | Cross-cont |
| *Bombycilla cedrorum* | Passeriformes | Bombycillidae | 0.5 | **69** | 0.37 | **74** | -- | **--** | -- | **--** | Yes | Cross-cont |
| *Passerina ciris* | Passeriformes | Cardinalidae | 1.71 | **97** | -- | **--** | -- | **--** | -- | **--** | -- | E and/or C |
| *Pheucticus ludovicianus* | Passeriformes | Cardinalidae | 1.13 | **89** | 0.6 | **79** | –0.28 | **40** | -- | **--** | Yes | Cross-cont |
| *Piranga olivacea* | Passeriformes | Cardinalidae | 1.04 | **87** | 0.19 | **69** | –0.16 | **55** | -- | **--** | Yes | E and/or C |
| *Piranga rubra* | Passeriformes | Cardinalidae | 0.57 | **75** | -- | **--** | –0.04 | **71** | -- | **--** | -- | Cross-cont |
| *Passerina cyanea* | Passeriformes | Cardinalidae | 0.34 | **60** | 1.17 | **87** | 0.31 | **81** | –1.15 | **15** | Yes | Cross-cont |
| *Cardinalis cardinalis* | Passeriformes | Cardinalidae | –0.18 | **40** | -- | **--** | -- | **--** | –0.66 | **28** | Yes | E and/or C |
| *Passerina caerulea* | Passeriformes | Cardinalidae | –1.65 | **5** | -- | **--** | 0.05 | **78** | -- | **--** | -- | Cross-cont |
| *Spiza americana* | Passeriformes | Cardinalidae | –1.51 | **7** | -- | **--** | -- | **--** | -- | **--** | -- | E and/or C |
| *Certhia americana* | Passeriformes | Certhiidae | 1.6 | **96** | 0.23 | **70** | 1.73 | **93** | 0.5 | **70** | Yes | Cross-cont |
| *Cyanocitta cristata* | Passeriformes | Corvidae | –0.52 | **26** | -- | **--** | -- | **--** | 0.11 | **55** | Yes | E and/or C |

| | | | | | | | | | | | | |
|---|---|---|---|---|---|---|---|---|---|---|---|---|
| *Corvus brachyrhynchos* | Passeriformes | Corvidae | –1.3 | **10** | -- | **--** | -- | **--** | -- | **--** | Yes | Cross-cont |
| *Pinicola enucleator* | Passeriformes | Fringillidae | 1.19 | **91** | -- | **--** | -- | **--** | -- | **--** | Yes | Cross-cont |
| *Carpodacus mexicanus* | Passeriformes | Fringillidae | 0.56 | **73** | -- | **--** | -- | **--** | -- | **--** | -- | Cross-cont |
| *Carduelis tristis* | Passeriformes | Fringillidae | 0.29 | **59** | –0.17 | **58** | -- | **--** | 0.75 | **82** | -- | Cross-cont |
| *Carpodacus purpureus* | Passeriformes | Fringillidae | 0.63 | **77** | –0.2 | **55** | -- | **--** | -- | **--** | Yes | Cross-cont |
| *Carduelis pinus* | Passeriformes | Fringillidae | 0.37 | **63** | –0.17 | **58** | -- | **--** | –0.48 | **32** | -- | Cross-cont |
| *Loxia leucoptera* | Passeriformes | Fringillidae | –0.34 | **32** | -- | **--** | -- | **--** | -- | **--** | Yes | Cross-cont |
| *Carduelis flammea* | Passeriformes | Fringillidae | –0.71 | **21** | -- | **--** | -- | **--** | -- | **--** | -- | Cross-cont |
| *Hirundo rustica* | Passeriformes | Hirundinidae | –1.53 | **6** | –0.27 | **50** | -- | **--** | -- | **--** | Yes | Cross-cont |
| *Stelgidopteryx serripennis* | Passeriformes | Hirundinidae | –1.42 | **9** | –0.39 | **36** | -- | **--** | -- | **--** | -- | Cross-cont |
| *Euphagus cyanocephalus* | Passeriformes | Icteridae | 0.06 | **51** | -- | **--** | -- | **--** | -- | **--** | Yes | Cross-cont |
| *Euphagus carolinus* | Passeriformes | Icteridae | –0.03 | **46** | -- | **--** | -- | **--** | -- | **--** | Yes | Cross-cont |
| *Icterus spurius* | Passeriformes | Icteridae | –0.16 | **41** | –0.04 | **63** | –0.35 | **35** | -- | **--** | -- | E and/or C |
| *Icterus galbula* | Passeriformes | Icteridae | –0.2 | **39** | –0.29 | **49** | –0.99 | **9** | -- | **--** | Yes | E and/or C |
| *Molothrus ater* | Passeriformes | Icteridae | –1.25 | **12** | –0.1 | **62** | -- | **--** | -- | **--** | Yes | Cross-cont |
| *Sturnella magna* | Passeriformes | Icteridae | –1.23 | **13** | –0.38 | **37** | –0.42 | **29** | -- | **--** | -- | E and/or C |
| *Quiscalus quiscula* | Passeriformes | Icteridae | –0.97 | **16** | –0.43 | **35** | -- | **--** | -- | **--** | Yes | E and/or C |
| *Agelaius phoeniceus* | Passeriformes | Icteridae | –1.98 | **4** | –0.7 | **22** | -- | **--** | -- | **--** | Yes | Cross-cont |
| *Dolichonyx oryzivorus* | Passeriformes | Icteridae | –2.83 | **1** | –0.43 | **35** | -- | **--** | -- | **--** | Yes | Cross-cont |
| *Icteria virens* | Passeriformes | Icteriidae | –0.06 | **45** | -- | **--** | –0.07 | **66** | -- | **--** | Yes | Cross-cont |
| *Dumetella carolinensis* | Passeriformes | Mimidae | 1.21 | **92** | 0.82 | **84** | –1.04 | **4** | 0.68 | **80** | Yes | Cross-cont |
| *Toxostoma rufum* | Passeriformes | Mimidae | 0.45 | **68** | –0.18 | **56** | –0.99 | **9** | 0.98 | **88** | Yes | E and/or C |

| | | | | | | | | | | | | |
|---|---|---|---|---|---|---|---|---|---|---|---|---|
| *Mimus polyglottos* | Passeriformes | Mimidae | –0.6 | **25** | -- | **--** | –0.15 | **60** | 0.2 | **60** | -- | Cross-cont |
| *Parus atricapillus* | Passeriformes | Paridae | 0.18 | **54** | -- | **--** | -- | **--** | -- | **--** | -- | Cross-cont |
| *Parus carolinensis* | Passeriformes | Paridae | –0.2 | **39** | -- | **--** | -- | **--** | -- | **--** | -- | E and/or C |
| *Baeolophus bicolor* | Passeriformes | Paridae | –0.38 | **29** | -- | **--** | -- | **--** | -- | **--** | -- | E and/or C |
| *Seiurus aurocapilla* | Passeriformes | Parulidae | 1.5 | **95** | 2.52 | **97** | 3.84 | **99** | 1.5 | **98** | -- | Cross-cont |
| *Dendroica caerulescens* | Passeriformes | Parulidae | 1.89 | **99** | -- | **--** | –0.19 | **48** | 2.86 | **100** | -- | E and/or C |
| *Vermivora peregrina* | Passeriformes | Parulidae | 0.57 | **75** | 3.01 | **99** | 1.93 | **96** | 0.14 | **57** | -- | Cross-cont |
| *Helmitheros vermivorum* | Passeriformes | Parulidae | 1.04 | **87** | -- | **--** | -- | **--** | -- | **--** | -- | E and/or C |
| *Geothlypis trichas* | Passeriformes | Parulidae | 1.13 | **89** | 1.92 | **95** | –0.18 | **51** | 0.67 | **78** | Yes | Cross-cont |
| *Vermivora ruficapilla* | Passeriformes | Parulidae | 0.67 | **79** | 1.26 | **88** | 0.79 | **88** | 0.53 | **72** | -- | Cross-cont |
| *Mniotilta varia* | Passeriformes | Parulidae | 1.14 | **90** | 1.51 | **91** | –0.33 | **36** | -- | **--** | Yes | Cross-cont |
| *Oporornis agilis* | Passeriformes | Parulidae | 1.52 | **96** | –0.3 | **46** | 0.68 | **87** | -- | **--** | Yes | Cross-cont |
| *Seiurus motacilla* | Passeriformes | Parulidae | 1.28 | **93** | -- | **--** | –0.06 | **69** | -- | **--** | -- | E and/or C |
| *Oporornis formosus* | Passeriformes | Parulidae | 1.03 | **85** | -- | **--** | 0.13 | **79** | -- | **--** | -- | E and/or C |
| *Dendroica dominica* | Passeriformes | Parulidae | 0.73 | **81** | -- | **--** | 0 | **74** | -- | **--** | -- | E and/or C |
| *Seiurus noveboracensis* | Passeriformes | Parulidae | 0.66 | **77** | 0.56 | **78** | –0.18 | **51** | -- | **--** | -- | Cross-cont |
| *Oporornis philadelphia* | Passeriformes | Parulidae | 0.79 | **84** | –0.21 | **52** | 0.05 | **78** | 0.48 | **68** | -- | Cross-cont |
| *Vermivora chrysoptera* | Passeriformes | Parulidae | 1.76 | **98** | –0.96 | **12** | –0.12 | **64** | -- | **--** | Yes | Western |
| *Dendroica pensylvanica* | Passeriformes | Parulidae | 0.6 | **76** | 0.48 | **77** | –0.48 | **27** | -- | **--** | -- | E and/or C |
| *Dendroica townsendi* | Passeriformes | Parulidae | 0.19 | **56** | -- | **--** | -- | **--** | -- | **--** | -- | Western |
| *Dendroica castanea* | Passeriformes | Parulidae | 0.71 | **80** | –0.29 | **49** | –0.04 | **71** | -- | **--** | -- | Cross-cont |
| *Wilsonia canadensis* | Passeriformes | Parulidae | 1.39 | **95** | –0.78 | **19** | –0.23 | **45** | -- | **--** | -- | E and/or C |

| | | | | | | | | | | | | |
|---|---|---|---|---|---|---|---|---|---|---|---|---|
| *Dendroica striata* | Passeriformes | Parulidae | 0.35 | **60** | 1.09 | **86** | –0.12 | **64** | –0.86 | **22** | -- | Cross-cont |
| *Dendroica tigrina* | Passeriformes | Parulidae | 1.1 | **87** | –0.32 | **40** | –0.52 | **26** | -- | **--** | -- | E and/or C |
| *Parula americana* | Passeriformes | Parulidae | 0.67 | **79** | –0.48 | **29** | –0.38 | **31** | 0.48 | **68** | -- | E and/or C |
| *Dendroica petechia* | Passeriformes | Parulidae | –0.21 | **37** | 1.45 | **90** | –1.06 | **2** | -- | **--** | -- | Cross-cont |
| *Dendroica cerulea* | Passeriformes | Parulidae | –0.08 | **43** | -- | **--** | –0.06 | **69** | -- | **--** | -- | E and/or C |
| *Setophaga ruticilla* | Passeriformes | Parulidae | 0.36 | **62** | 0.35 | **73** | –0.68 | **21** | –0.36 | **38** | -- | Cross-cont |
| *Dendroica magnolia* | Passeriformes | Parulidae | 0.41 | **65** | –0.68 | **23** | –0.15 | **60** | 0.04 | **50** | -- | Cross-cont |
| *Dendroica virens* | Passeriformes | Parulidae | 0.37 | **63** | –1.03 | **10** | –0.46 | **28** | 0.65 | **75** | -- | E and/or C |
| *Dendroica pinus* | Passeriformes | Parulidae | –0.12 | **43** | –0.3 | **46** | –0.14 | **61** | -- | **--** | -- | E and/or C |
| *Dendroica fusca* | Passeriformes | Parulidae | 0.31 | **59** | –0.67 | **24** | –0.26 | **43** | -- | **--** | -- | E and/or C |
| *Vermivora pinus* | Passeriformes | Parulidae | 0.76 | **82** | –1.45 | **3** | –0.15 | **60** | -- | **--** | -- | E and/or C |
| *Wilsonia citrina* | Passeriformes | Parulidae | –0.62 | **24** | -- | **--** | –0.23 | **45** | -- | **--** | -- | E and/or C |
| *Dendroica palmarum* | Passeriformes | Parulidae | 0.01 | **51** | –0.57 | **26** | –0.89 | **13** | -- | **--** | -- | E and/or C |
| *Wilsonia pusilla* | Passeriformes | Parulidae | –0.32 | **32** | –0.37 | **38** | –0.85 | **16** | –0.97 | **18** | -- | Cross-cont |
| *Vermivora celata* | Passeriformes | Parulidae | –1.04 | **14** | –0.73 | **20** | –0.37 | **33** | -- | **--** | -- | Cross-cont |
| *Protonotaria citrea* | Passeriformes | Parulidae | –1.84 | **5** | –0.53 | **27** | –0.1 | **65** | -- | **--** | -- | E and/or C |
| *Dendroica coronata* | Passeriformes | Parulidae | –0.43 | **27** | –0.82 | **18** | –0.9 | **12** | –1.87 | **2** | -- | Cross-cont |
| *Zonotrichia albicollis* | Passeriformes | Passerellidae | 0.63 | **77** | 3.1 | **100** | 4.5 | **100** | –0.41 | **35** | Yes | Cross-cont |
| *Passerella iliaca* | Passeriformes | Passerellidae | 0.53 | **71** | -- | **--** | 1.84 | **94** | -- | **--** | Yes | Cross-cont |
| *Junco hyemalis* | Passeriformes | Passerellidae | –0.05 | **46** | 2.18 | **96** | 2.96 | **98** | –0.75 | **25** | Yes | Cross-cont |
| *Zonotrichia atricapilla* | Passeriformes | Passerellidae | 1.04 | **87** | -- | **--** | -- | **--** | -- | **--** | -- | Western |
| *Melospiza georgiana* | Passeriformes | Passerellidae | 1.35 | **94** | 0.38 | **75** | 2.47 | **97** | –0.14 | **45** | Yes | Cross-cont |

| | | | | | | | | | | | | |
|---|---|---|---|---|---|---|---|---|---|---|---|---|
| *Melospiza melodia* | Passeriformes | Passerellidae | 0.39 | **64** | –0.16 | **60** | 1.39 | **91** | -- | **--** | Yes | Cross-cont |
| *Melospiza lincolnii* | Passeriformes | Passerellidae | 0.53 | **71** | 0.32 | **72** | 0.92 | **90** | –0.32 | **42** | Yes | Cross-cont |
| *Spizella arborea* | Passeriformes | Passerellidae | 0.13 | **52** | -- | **--** | 0.57 | **84** | -- | **--** | -- | Cross-cont |
| *Spizella pallida* | Passeriformes | Passerellidae | 0.1 | **52** | 0.76 | **83** | –0.27 | **42** | 0.06 | **52** | Yes | E and/or C |
| *Ammodramus henslowii* | Passeriformes | Passerellidae | 0.2 | **57** | –0.29 | **49** | 0.02 | **75** | -- | **--** | -- | E and/or C |
| *Ammodramus savannarum* | Passeriformes | Passerellidae | 0.55 | **71** | –1.37 | **6** | 0.41 | **82** | -- | **--** | Yes | E and/or C |
| *Spizella passerina* | Passeriformes | Passerellidae | 0.01 | **51** | 1.01 | **85** | –0.75 | **19** | –1.42 | **10** | Yes | Cross-cont |
| *Zonotrichia leucophrys* | Passeriformes | Passerellidae | –0.68 | **23** | -- | **--** | –0.57 | **24** | -- | **--** | Yes | Cross-cont |
| *Ammodramus leconteii* | Passeriformes | Passerellidae | –1.41 | **9** | -- | **--** | –0.02 | **72** | -- | **--** | -- | E and/or C |
| *Pipilo erythrophthalmus* | Passeriformes | Passerellidae | 0.01 | **51** | –1.38 | **5** | –0.9 | **12** | -- | **--** | Yes | E and/or C |
| *Spizella pusilla* | Passeriformes | Passerellidae | 0.16 | **54** | –2.41 | **1** | –0.36 | **34** | -- | **--** | -- | E and/or C |
| *Passerculus sandwichensis* | Passeriformes | Passerellidae | –1.42 | **9** | –1.12 | **8** | –0.68 | **21** | -- | **--** | Yes | Cross-cont |
| *Pooecetes gramineus* | Passeriformes | Passerellidae | –3.18 | **1** | –1.42 | **4** | –0.15 | **60** | -- | **--** | Yes | Cross-cont |
| *Passer domesticus* | Passeriformes | Passeridae | –0.34 | **32** | -- | **--** | -- | **--** | –0.49 | **30** | Yes | Cross-cont |
| *Polioptila caerulea* | Passeriformes | Polioptilidae | –2.24 | **4** | –0.46 | **32** | –1.04 | **4** | -- | **--** | -- | Cross-cont |
| *Regulus satrapa* | Passeriformes | Regulidae | 0.57 | **75** | –0.82 | **18** | –0.2 | **47** | -- | **--** | -- | Cross-cont |
| *Regulus calendula* | Passeriformes | Regulidae | –0.01 | **49** | 0.74 | **81** | –1.13 | **1** | –1.5 | **8** | -- | Cross-cont |
| *Sitta pygmaea* | Passeriformes | Sittidae | 1.15 | **90** | -- | **--** | -- | **--** | -- | **--** | -- | Western |
| *Sitta carolinensis* | Passeriformes | Sittidae | 0.4 | **65** | 1.66 | **93** | -- | **--** | 0.86 | **85** | Yes | Cross-cont |
| *Sitta canadensis* | Passeriformes | Sittidae | 0.45 | **68** | –0.91 | **13** | –0.13 | **62** | -- | **--** | Yes | Cross-cont |
| *Sturnus vulgaris* | Passeriformes | Sturnidae | –0.85 | **18** | -- | **--** | -- | **--** | -- | **--** | Yes | Cross-cont |
| *Troglodytes aedon* | Passeriformes | Troglodytidae | –0.21 | **37** | 0.14 | **68** | –0.78 | **18** | -- | **--** | Yes | Cross-cont |

| | | | | | | | | | | | | |
|---|---|---|---|---|---|---|---|---|---|---|---|---|
| *Troglodytes troglodytes* | Passeriformes | Troglodytidae | –0.23 | **36** | –0.47 | **30** | –0.17 | **53** | -- | -- | -- | E and/or C |
| *Cistothorus platensis* | Passeriformes | Troglodytidae | –0.38 | **29** | -- | -- | -- | -- | -- | -- | -- | E and/or C |
| *Cistothorus palustris* | Passeriformes | Troglodytidae | –0.37 | **30** | –0.83 | **16** | –0.28 | **40** | -- | -- | -- | Cross-cont |
| *Thryothorus ludovicianus* | Passeriformes | Troglodytidae | –0.86 | **18** | –0.14 | **62** | -- | -- | -- | -- | -- | E and/or C |
| *Myadestes townsendi* | Passeriformes | Turdidae | 1.77 | **98** | -- | -- | -- | -- | -- | -- | -- | Western |
| *Hylocichla mustelina* | Passeriformes | Turdidae | 0.71 | **80** | –1.16 | **7** | 0.45 | **83** | 1.05 | **90** | Yes | E and/or C |
| *Catharus minimus* | Passeriformes | Turdidae | 0.25 | **57** | –0.25 | **51** | 0.63 | **85** | 0.42 | **62** | Yes | Cross-cont |
| *Catharus guttatus* | Passeriformes | Turdidae | 0.36 | **62** | –0.85 | **15** | 1.56 | **92** | –0.06 | **48** | Yes | Cross-cont |
| *Catharus ustulatus* | Passeriformes | Turdidae | 0.14 | **53** | 0.12 | **67** | 0.84 | **89** | –0.34 | **40** | Yes | Cross-cont |
| *Catharus fuscescens* | Passeriformes | Turdidae | 0.26 | **58** | –0.46 | **32** | 0.27 | **80** | -- | -- | Yes | Cross-cont |
| *Turdus migratorius* | Passeriformes | Turdidae | –0.49 | **27** | 1.9 | **94** | -- | -- | –1.82 | **5** | Yes | Cross-cont |
| *Sialia sialis* | Passeriformes | Turdidae | –1.28 | **11** | 0.69 | **80** | -- | -- | -- | -- | -- | E and/or C |
| *Zoothera naevia* | Passeriformes | Turdidae | –0.37 | **30** | -- | -- | -- | -- | -- | -- | -- | Western |
| *Empidonax minimus* | Passeriformes | Tyrannidae | –0.28 | **35** | 1.31 | **89** | –1 | **7** | -- | -- | Yes | Cross-cont |
| *Contopus virens* | Passeriformes | Tyrannidae | –0.02 | **47** | 0.75 | **82** | –0.8 | **17** | -- | -- | Yes | E and/or C |
| *Empidonax virescens* | Passeriformes | Tyrannidae | –0.62 | **24** | -- | -- | 0 | **74** | -- | -- | -- | E and/or C |
| *Contopus cooperi* | Passeriformes | Tyrannidae | –0.12 | **43** | –0.6 | **25** | –0.22 | **46** | -- | -- | -- | Cross-cont |
| *Empidonax flaviventris* | Passeriformes | Tyrannidae | –0.31 | **33** | –0.36 | **38** | –0.38 | **31** | -- | -- | Yes | Cross-cont |
| *Myiarchus crinitus* | Passeriformes | Tyrannidae | –0.29 | **34** | –1.08 | **9** | –0.53 | **25** | -- | -- | -- | E and/or C |
| *Sayornis phoebe* | Passeriformes | Tyrannidae | –0.78 | **20** | –0.51 | **28** | –0.94 | **10** | -- | -- | Yes | E and/or C |
| *Tyrannus tyrannus* | Passeriformes | Tyrannidae | –1.27 | **12** | –0.31 | **43** | –0.85 | **16** | -- | -- | -- | Cross-cont |
| *Empidonax alnorum* | Passeriformes | Tyrannidae | –2.31 | **3** | –0.86 | **14** | -- | -- | -- | -- | Yes | Cross-cont |

| | | | | | | | | | | | |
|---|---|---|---|---|---|---|---|---|---|---|---|
| *Vireo solitarius* | Passeriformes | Vireonidae | –0.16 | **41** | –0.71 | **21** | –0.3 | **38** | -- | **--** | Yes | E and/or C |
| *Vireo griseus* | Passeriformes | Vireonidae | –0.78 | **20** | -- | **--** | –0.17 | **53** | -- | **--** | -- | E and/or C |
| *Vireo flavifrons* | Passeriformes | Vireonidae | –0.28 | **35** | –1.02 | **11** | –0.16 | **55** | -- | **--** | -- | E and/or C |
| *Vireo olivaceus* | Passeriformes | Vireonidae | –1.01 | **15** | 0.1 | **66** | –0.63 | **22** | –1.29 | **12** | Yes | Cross-cont |
| *Vireo philadelphicus* | Passeriformes | Vireonidae | –1.51 | **7** | -- | **--** | –0.3 | **38** | -- | **--** | Yes | Cross-cont |
| *Vireo gilvus* | Passeriformes | Vireonidae | –1.32 | **10** | –0.92 | **12** | –1.01 | **6** | -- | **--** | -- | Cross-cont |
| *Ixobrychus exilis* | Pelecaniformes | Ardeidae | –0.6 | **25** | -- | **--** | -- | **--** | -- | **--** | Yes | E and/or C |
| *Botaurus lentiginosus* | Pelecaniformes | Ardeidae | –2.32 | **2** | -- | **--** | -- | **--** | -- | **--** | Yes | Cross-cont |
| *Sphyrapicus varius* | Piciformes | Picidae | 1.34 | **93** | 1.52 | **92** | -- | **--** | 1.35 | **92** | Yes | Cross-cont |
| *Colaptes auratus* | Piciformes | Picidae | 0.42 | **66** | 0.02 | **64** | -- | **--** | -- | **--** | Yes | Cross-cont |
| *Picoides pubescens* | Piciformes | Picidae | –0.01 | **49** | -- | **--** | -- | **--** | -- | **--** | -- | Cross-cont |
| *Melanerpes erythrocephalus* | Piciformes | Picidae | –0.54 | **26** | –0.14 | **62** | -- | **--** | -- | **--** | Yes | E and/or C |
| *Picoides villosus* | Piciformes | Picidae | –0.7 | **21** | -- | **--** | -- | **--** | -- | **--** | -- | Cross-cont |
| *Melanerpes carolinus* | Piciformes | Picidae | –0.36 | **30** | –1.5 | **2** | -- | **--** | -- | **--** | -- | E and/or C |
| *Podilymbus podiceps* | Podicipediformes | Podicipedidae | 1.13 | **89** | -- | **--** | -- | **--** | -- | **--** | Yes | Cross-cont |
| *Asio otus* | Strigiformes | Strigidae | 0.19 | **56** | -- | **--** | -- | **--** | -- | **--** | Yes | Cross-cont |
| *Aegolius acadicus* | Strigiformes | Strigidae | –0.24 | **35** | -- | **--** | -- | **--** | -- | **--** | Yes | Cross-cont |
